# Relationship as resource or burden? Associations of attachment style, relationship quality and dyadic coping with acute psychosocial stress in the presence of the romantic partner

**DOI:** 10.1101/2024.10.21.618832

**Authors:** Mathilde Gallistl, Johanna Hamann, Ilona Croy, Pascal Vrticka, Veronika Engert

**Author notes:** Corresponding author: Mathilde Gallistl Max Planck Institute for Human Cognitive and Brain Sciences Social Stress and Family Health Research Group Stephanstr. 1A, 04103 Leipzig, Germany. joint first authors (Mathilde Gallistl, Johanna Hamann). joint last authors (Pascal Vrticka, Veronika Engert).

## Abstract

Stress is a wide-spread phenomenon and associated with various detrimental health effects. A significant resource for stress buffering is social support. How social support is perceived, however, depends on a multitude of individual and interindividual factors. This study aimed to explore the stress-reducing properties of relationship-inherent variables. We investigated the association of attachment style, relationship quality and dyadic coping, with subjective and physiological stress responses to a psychosocial laboratory stressor in romantic partners. Seventy-nine couples participated, with one partner ("target") undergoing the Trier Social Stress Test and the other ("observer") observing the situation. Besides examining the role of targets’ relationship variables, we also assessed the link between observers’ relationship variables and targets’ stress reactivity. We found that both targets’ and observers’ insecure-avoidant attachment scores were associated with targets’ stress reactivity. In detail, while targets’ insecure-avoidant attachment scores were negatively associated with targets’ subjective stress experience, observers’ insecure-avoidant attachment scores were positively associated with targets’ heart rate reactivity. Further, higher insecure-avoidant attachment scores linked to lower psycho-endocrine covariance, i.e., a lower accordance between self-reported and cortisol stress responding. On the one hand, these data may suggest that under stress, insecure-avoidantly attached individuals suppress their experience of stress to preserve a sense of independence as part of their deactivating attachment strategy. The presence of an insecure-avoidantly attached partner during a stressful experience, on the other hand, seems to be a stressor rather than a source of support. Long-term, an insecure-avoidantly attached partner may negatively impact an individual’s stress-related health and wellbeing.

**Highlights:** - In a situation in which one romantic partner experienced stress (“target”) while being ob- served by their partner (“observer”), insecure-avoidant attachment of both predicted the tar- get’s stress response

- Targets’ insecure-avoidant attachment was associated with lower subjective stress in the tar- gets

- Observers’ insecure-avoidant attachment was related to higher heart rate reactivity in the targets

- Insecure-avoidant attachment was associated with lower psycho-endocrine covariance

## 1. Introduction

Stress is an omnipresent phenomenon in modern Western societies. While some degree of stress represents an adaptive response to acute external and internal demands (Dhabhar, 2018), chronically elevated stress levels have detrimental effects on mental and physical health (Chrousos, 2009; McEwen, 2008; Sapolsky, 2004). A significant resource for buffering stress is social support (DeLongis & Holtzman, 2005; Sidney, 1976; Thoits, 2010; White et al., 2023), especially if provided by one’s spouse (Dean et al., 1990). Social support is the perception that one is cared for, has assistance available, and is part of a supportive network. This can include emotional support, informational guidance, or a sense of belonging (Drageset, 2021), for instance through the companionship of a partner being present. However, the extent to which a close relationship qualifies as a resource in stressful situations depends on multiple individual and interindividual factors. Amongst these are relationship quality (DeLongis & Holtzman, 2005) communication (Ditzen et al., 2008) and attachment patterns (Fuenfhausen & Cashwell, 2013; White et al., 2023). To gain a better understanding of some of the factors involved in the stress-regulating properties of social support in acutely challenging situations, we subjected participants (here termed “targets”) to a standardized psychosocial laboratory stressor, the Trier Social Stress Test (TSST; Kirschbaum et al., 1993), while their romantic partners passively attended the stressful situation (here termed “observers”). The strength of the targets’ stress response (measured in terms of cortisol, heart rate, heart rate variability, and self-reported stress reactivity) was then related to their partners’ and their own attachment style as well as to two other relationship variables of relationship quality and dyadic coping. Our aim was to identify those relationship-related factors that were associated with most effective target stress coping in the presence of their partner.

Attachment is one factor that can influence targets’ perception of their partners’ social support during stress, and particularly whether the provided support is seen as available and helpful for buffering the impact of the experienced stress (White et al., 2023). In adulthood, attachment styles are best understood as predictions about others’ availability and one’s own capacity to elicit help from others when needed, and is mostly classified as secure, insecure-anxious and insecure-avoidant (Hazan & Shaver, 1987). Securely attached individuals can effectively cope with distress, while insecurely attached individuals are characterized by less optimal stress regulation (Mikulincer & Shaver, 2010). More specifically, insecure-anxious attachment is characterized by chronic worries about others’ availability and rejection. These worries lead to an intensification of coping strategies involving heightened support-seeking, reflecting an overall hyperactivation of the attachment system. On the other hand, insecure-avoidant attachment is characterized by coping strategies aiming to maximize independence and self-reliance. These strategies involve the suppression of negative thoughts and relationship-related feelings, which are consistent with an overall hypoactivation of the attachment system (Simpson & Rholes, 2017). The attachment system is known to be most likely and strongly activated by acute stressors (Mikulincer & Shaver, 2003), with insecure attachment previously being linked to higher acute stress responding (e.g. Beck et al., 2013; Pierrehumbert et al., 2012). Furthermore, in the context of relationship-related stressors, insecure attachment was shown to be associated with higher cortisol release (e.g. Dewitte et al., 2010; Laurent & Powers, 2007) and skin conductance (associated with sympathetic nervous activity; Diamond et al., 2006).

Previous studies investigating attachment-related associations between social support and acute psychosocial stress reactivity found that for insecure-avoidantly attached individuals, the presence of their romantic partner during a stressful experience was associated with increased heart rate and systolic blood pressure activation (Carpenter & Kirkpatrick, 1996). Others have shown that after insecure-avoidantly attached individuals had experienced a stressor alone, they showed increased psychophysiological activation upon reunion with their partner (Meuwly et al., 2012). Taking the reverse perspective, Dewitte et al. (2010) found that during a relational distress paradigm observing partners’ attachment style was associated with targets’ stress reactivity, with partners of securely attached targets exhibiting lowest cortisol reactivity.

The nature of the relationship also plays an important role when undergoing stressful situations together. For instance, whether a partner is seen as a helpful resource is linked to the perception of that partner’s degree of support. During acute stress, social support in terms of dyadic coping, which involves partners working together to manage stress (Bodenmann, 2008), was linked to better stress recovery after the TSST (Meuwly et al., 2012). More generally, it is known that relationship quality is a strong predictor of dyadic coping (Falconier et al., 2015; Herzberg, 2013; Rusu et al., 2020) and to be closely tied to attachment (Banse, 2004; Cann et al., 2008; Fuenfhausen & Cashwell, 2013; Meuwly et al., 2012) as well as general stress sensitivity (Fuenfhausen & Cashwell, 2013; Otis et al., 2006). Despite these connections, the role of relationship quality in jointly experienced stress remains virtually unstudied. One single study employing the TSST found a negative correlation between relationship quality and subjective stress in women, but not in men, and this stress paradigm did not involve partner presence (Meuwly et al., 2012).

In the current study, we tested 79 opposite-sex romantic partner dyads to investigate the associations of self-reported attachment, dyadic coping, and relationship quality of both partners on targets’ stress reactivity measured via cortisol, heart rate, heart rate variability and subjective stress during the TSST. Social support was operationalized as the presence of the partner during the stress test. Based on the literature reviewed above and on conceptualizations of attachment theory, we expected overall insecure attachment of either targets or observers to be associated with higher target stress. Furthermore, solely relatively increased target insecure-avoidant attachment was expected to be associated with lower target subjective stress. We further hypothesized to find lower target stress reactivity with higher target- or observer-related relationship quality and dyadic coping.

## 2. Methods

**2.1 Participants**

As part of a larger research project investigating empathic processes in romantic partners during stress, a total of 85 opposite-sex couples (170 individuals) were invited to the Max Planck Institute for Human Cognitive and Brain Sciences in Leipzig, Germany. Six of the 85 dyads dropped out before the stress testing session, leaving 79 dyads to be included in the current analysis. Participants were between 19 and 38 years of age (*M* = 26.03, *SD* = 4.42), most of them students or working in academic careers (78.2%). Smokers (Rohleder & Kirschbaum, 2006), regular recreational drug users (Parrott, 2015), individuals having received psychotherapeutic treatment for an ICD-10 diagnosis over the last two years, or individuals suffering from chronic physical diseases (Knezevic et al., 2023) were excluded due to possible cofounding effects on the physiological stress response. Likewise, individuals taking medications affecting cortisol regulation (Granger et al., 2009) and women on hormonal contraceptives (Hertel et al., 2017) were excluded. Given that cortisol levels are significantly affected by sex hormones (Kajantie & Phillips, 2006), the study included only naturally cycling women in their luteal phase. Also, because measures of brain activity were collected within the context of the larger study, left-handed individuals were excluded. All participants gave written informed consent, were financially compensated for their time and effort, and could withdraw from study participation at any time.

### 2.2 Experimental design and procedure

Participants attended two testing sessions on separate days. The first session lasted approximately four hours. On that day, one of the participants (the target; *n* = 40 male and *n* = 39 female) completed the Trier Social Stress Test (TSST; Kirschbaum et al., 1993), while their partner (the observer) was present in the same room and watched the situation. Before and after the TSST, targets and observers rested in separate rooms and repeatedly completed questionnaires assessing acute feelings of stress. Over the course of the entire testing session, ten saliva samples were collected to determine levels of cortisol release, and an electrocardiogram (ECG) was recorded over a period of approximately 115 minutes to gauge sympathetic and parasympathetic nervous system activity. Throughout the TSST, including a brain activity “baseline” task during which targets read a story aloud for five minutes, targets’ and observers’ brain activity was measured using functional near-infrared spectroscopy (fNIRS). These data are not subject to the current analysis (publications currently in preparation). For a depiction of the first testing session, see Figure 1. After the TSST, observers and targets returned to their respective waiting rooms for a recovery phase of 60 minutes, during which they completed several trait questionnaires assessing attachment styles, relationship quality, and dyadic coping. Although the experimental setup also provided a large volume of data after the TSST (i.e., during the recovery phase), we focused on the actual stress experience during the TSST (i.e., stress reactivity) because partners were together only during this time.

**Figure 1:**
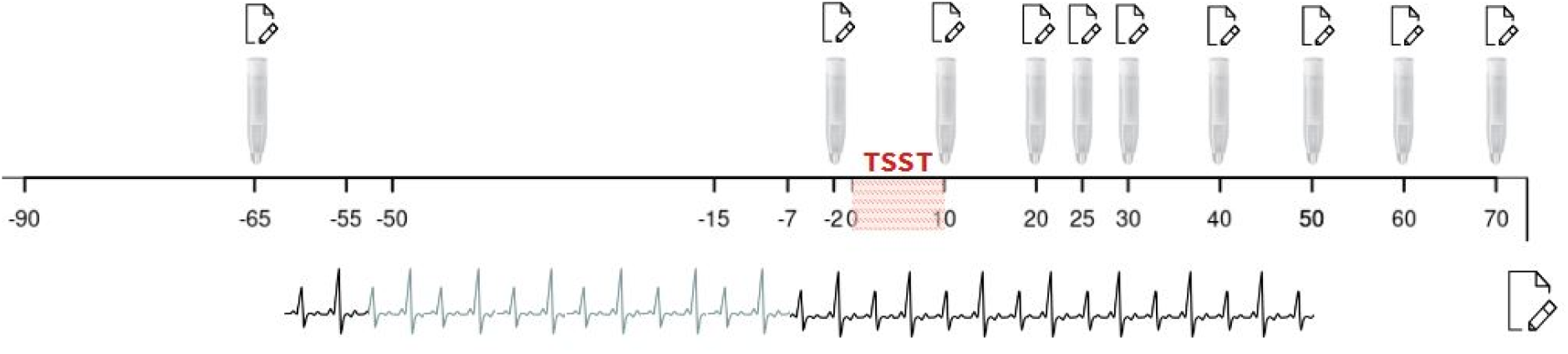
Timeline of the first testing session. *Note.* Numbers on the timeline indicate minutes relative to the stressor onset. The testing session started at -90 minutes prior to the TSST. The five minutes stress anticipation phase started at -7 minutes. TSST stress induction started at 0 minutes.

As part of the second testing session, participants’ empathic processing was measured using a 45-minute video task, the EmpaToM (Kanske et al., 2015). Subsequently, the Adult Attachment Interview (AAI; George et al., 1985) was conducted with the observers only. The AAI is a 45-90 minute structured interview that uses attachment-activating questions to elicit the biographical background with primary attachment figures to classify attachment representations. Because this measure was only implemented in the observing partners due to time and economic restrictions, it was not considered in the current data analysis. Both the EmpaToM and AAI data are reported elsewhere (Gallistl et al., 2024; Gallistl et al., 2024 in review).

### 2.3 Stress induction

Stress was induced using the Trier Social Stress Test (TSST; Kirschbaum et al., 1993) a standardized psychosocial stress paradigm that reliably activates the HPA axis and sympathetic nervous system (Dickerson & Kemeny, 2004). The TSST comprises three five-minute phases. The first phase consists of a preparatory stress anticipation phase. This is followed by second phase comprising a free speech task in which targets are asked to complete a mock job interview. In the third phase, participants must then complete a difficult mental arithmetic task. During the entire duration of the TSST, targets are under the impression that they are being evaluated by two committee members, presented as trained behavioral analysts and instructed not to give any feedback. Targets’ performance is also audio- and videotaped. In our study, observers were seated in the TSST testing room directly facing targets. Targets were instructed to only look at the committee, observers were asked to watch only the target. Observers had no task besides passively witnessing the situation. Due to the concomitantly acquired fNIRS recordings, targets and observers were asked to sit rather than stand during the TSST.

### 2.4 Measures

#### 2.4.1 Attachment

Target and observer attachment was measured using a validated German version of the Experience in Close Relationship Scale-Revised (ECR-RD; Ehrenthal et al., 2009), which is known to be well suited to dimensionally assess attachment (Ravitz et al., 2010). This questionnaire includes 36 items, 18 for each of the two attachment dimensions of anxiety (e.g. “I’m afraid that I will lose my partner’s love.”) and avoidance (e.g. “I am nervous when partners get too close to me.”). All items are answered on a continuous 7-point scale (ranging from 1, “do not agree at all” to 7, “fully agree”). With Cronbach’s alpha of .84 for anxiety and .85 for avoidance, the ECR-RD demonstrated good internal consistency in our study.

#### 2.4.2 Dyadic Coping and Relationship Quality

To survey dyadic coping, the Dyadic Coping Inventory (DCI; Bodenmann, 2008) was used. Composed of five subscales, the DCI assesses joint stress management within romantic relationships, as well as individuals’ satisfaction with joint stress management. The five subscales measure own supporting behavior (e.g. “I show empathy and understanding to my partner”), own stress communication (e.g. “I tell my partner openly how I feel and that I would appreciate his/her support”), partners’ supporting behavior (e.g. “My partner shows empathy and understanding to me”), partners’ stress communication (e.g. “My partner tells me openly how he/she feels and that he/she would appreciate support”), and joint stress managing behavior (e.g. “We try to cope with the problem together and search for ascertained solutions”). The DCI is answered on a continuous scale from 1 (“very rarely” to 5 (“very often”). For the current analyses, the DCI total score was used. It showed a good Cronbach’s alpha of .89. Relationship quality was measured with the German version (Sander & Böcker, 1993) of the Relationship Assessment Scale (RAS; Hendrick, 1988). The RAS consists of seven items (e.g. "How well does your partner meet your needs?") answered on a 1 (“do not agree”) to 5 (“strongly agree”) continuous scale and had a good Cronbach’s alpha of .87.

#### 2.4.3 Subjective stress

To measure acute subjective stress in targets and observers, the 20-item state scale of the State-Trait Anxiety Inventory (STAI; Spielberger et al., 1970) was used in a German translation by Laux and colleagues (Laux et al., 1981). Participants answered self-describing statements (e.g. "I am relaxed" or "I am worried") on a continuous scale from 1 (“almost never”) to 4 (“almost always”). The STAI was completed as a pencil paper questionnaire at ten time points before, during and after the TSST (-50, -2, 10, 20, 25, 30, 40, 50, 60, 70 min relative to stressor onset at 0 min; see Figure 1).

#### 2.4.4 Salivary cortisol

Cortisol levels to gauge HPA axis activity were determined from saliva also collected at 10 measurement time points (-50, -2, 10, 20, 25, 30, 40, 50, 60, 70 min relative to stressor onset at 0 min; see Figure 1). Saliva was collected using Salivette devices (Sarstedt, Nümbrecht, Germany), which consist of a collection swab in a plastic container. Participants placed the collection swab in their mouth for 2 min and refrained from chewing. To ensure that samples were not contaminated with external particles, participants did not eat or drink anything other than water during the entire testing session. To equalize blood sugar levels, participants were given a snack upon arrival at the institute. Salivettes were stored at -20°C until analysis at the biochemical laboratory of the Department of Biological and Clinical Psychology at Trier University. Cortisol concentrations were measured with a time-resolved fluorescence immunoassay, exhibiting intra- and inter-assay variability of less than 10% and 12%, respectively (Dressendörfer et al., 1992).

#### 2.4.5 Heart rate and heart rate variability

Heart rate and heart rate variability were derived from a continuous ECG recorded with a portable ECG-device, the Zephyr Bioharness 3 chest belt, which was placed around targets’ chests at the beginning of the testing session (-65 min). It was removed 50 min after the start of the TSST, resulting in a measurement time-frame of 115 min. A continuous RR-interval tachogram was automatically created and manually cleaned from low quality data and artifacts, both using an in-house software. Heart rate was derived from that RR-interval tachogram, defined as reciprocal of the RR-interval in units of beats per minute. High frequency heart rate variability (HF-HRV) was then calculated with the hrv-analysis python package (Champseix et al., 2021). For the current analysis, heart rate and heart rate variability averages were extracted for the relevant phases of the testing session, that is, the baseline phase (initial 10 minutes of recording; from -65 to -55 minutes; Figure 1), as well as each phase of the TSST (5 minutes for anticipation, speech and math task each).

### 2.5 Statistical Analysis

Data preparation and all analyses were computed using R version 4.2.2 (R Core Team, 2022). Some modifications were applied to the dataset prior to conducting the analyses. To control for a lack of normal distribution, cortisol, heart rate and heart rate variability data was log-transformed.

In addition, to appropriately control for the possible influence of extreme values, any outliers were winzorized to three standard deviations above or below the mean. Further, participant age, attachment anxiety and avoidance as well as dyadic coping and relationship quality were z-standardized.

#### 2.5.1 Modulatory factors of stress

To investigate the association between target and observer attachment, dyadic coping and relationship quality with targets’ stress response, we calculated eight linear models in total. Four linear models were calculated with targets’ acute stress reactivity in subjective stress (1), cortisol (2), heart rate (3) and heart rate variability (4) as dependent variables, and *targets’* insecure-anxious and -avoidant attachment scores, dyadic coping and relationship quality as predictors in each of the models. Another four linear models were calculated again with targets’ stress reactivity in subjective stress (5), cortisol (6), heart rate (7) and heart rate variability (8) as dependent variables, but with *observers’* insecure-anxious and -avoidant attachment scores, dyadic coping and relationship quality as predictors in each of the models.

To gauge acute stress reactivity, change scores between targets’ stress peak and baseline values were calculated for each of the four stress markers. The stress baseline value was represented by the first STAI measurement (at -50min) for subjective stress, the lowest value of the first two cortisol measurements, as well as the first ten minutes (baseline phase) of the recorded ECG data for heart rate and heart rate variability. For the stress peak, the average peak timepoint for each of the four stress markers was calculated. Individual values at that peak timepoint represented the stress peak for each target. To minimize the influence of baseline levels on stress reactivity, reactivity change scores of all physiological markers were adjusted for the respective baseline (see: Wilder’s law of initial value; Benjamin, 1963; Wilder, 1962). This was done by extracting standardized residuals of a regression model predicting the change scores from baseline values.

With targets’ age showing high correlations with all predictors, it was included as a covariate in the eight models. In the prediction of cortisol, target sex and time of day were included as covariates due to their known influence on cortisol levels (Kudielka et al., 2004; Uhart et al., 2006). Targets’ sex (Antelmi et al., 2004) and Body Mass Index (BMI, Molfino et al., 2009) were included in the heart rate and HF-HRV models. Since two analyses were calculated for each dependent variable (one for *targets’* and for *observers’* perspective), the level of significance was Bonferroni-corrected to .025. For heart rate and HF-HRV, it was further reduced to .0125 because the two variables belong to the same stress system.

#### 2.5.2. Analysis of psychoendocrine covariance

Psychoendocrine covariance is the correspondence between subjective psychological and endocrine stress markers. In stress research, we typically see a lack of this covariance (for a meta-analysis, see Campbell & Ehlert, 2012), likely due to known biases in self-report data (e.g. social desirability, extreme responding). This was found particularly in individuals who score higher on insecure-avoidant attachment (Diamond et al., 2006), and is linked to insecure-avoidantly attached individuals’ tendencies to cope with stressors by suppressing negative feelings and thereby signaling independence to the outside world (Simpson & Rholes, 2017). Accordingly, we expected in this study that insecure-avoidantly attached targets would select lower stress scores on the subjective stress scale and try not to signal their distress to their partners (i.e., observers). To examine this assumption, the relationship between psychoendocrine covariance and attachment scores was assessed in all individuals (both targets and observers). For this purpose, we took the STAI values (psychological stress) at six time points (-2, 10, 20, 30, 40, 50 minutes) and matched these values with chronologically corresponding cortisol values (physiological stress). This was done by calculating the difference of the time onset between the STAI and cortisol peak measurements in all participants, which was 25 minutes. Since cortisol measurement ended with a general time difference of two minutes relative to the STAI measurement onset (thus 23 minutes time difference remaining), and it led to more times points to be matched due to the experimental design, we decided on matching STAI and cortisol measurements with a time difference of 20 minutes (i.e. cortisol measurements at time points 20, 30, 40, 50, 60 and 70 minutes). A correlation value between psychological and physiological stress was calculated for each target and observer individually across these six time points. This correlation value (representing psychoendocrine covariance) was subsequently entered into a linear model as the dependent variable, with insecure- anxious and -avoidant attachment scores as independent variables, controlled for age, sex and time of day.

## 3. Results

Means, standard deviations and correlations between all variables are reported in the Supplementary Material. Unpaired two-sample t-tests showed no differences between targets and observers in the four predictor variables (insecure-anxious attachment: *t*(155) = 0.36, *p* = 0.72; insecure-avoidant attachment: *t*(143.4) = 0.45, *p* = 0.65; relationship quality: *t*(153.87) = -0.05, *p* = 0.96; dyadic coping: *t*(155) = 0.36, *p* = 0.72). There were no significant sex differences in scores of insecure-anxious attachment (*t*(152.87) = -0.49, *p* = 0.63), insecure-avoidant attachment (*t*(153.81) = -0.19, *p* = 0.85), relationship quality (*t*(154.91) = 0.20, *p* = 0.84), or dyadic coping (*t*(154.99) = 1.24, *p* = 0.22). The sample of N=79 dyads was reduced to 66 for autonomous data due technical problems with the heart rate registration devices. There were no reductions in the other stress markers due to systematic reasons.

### 3.1 Stress modulating factors

As shown in Figure 2, targets responded to the TSST on all stress markers. 84,8% of targets (67 out of 79) reached the accepted criterion for physiologically significant cortisol release (>1.5nmol/l above baseline levels; Miller et al., 2013).

**Figure 2:**
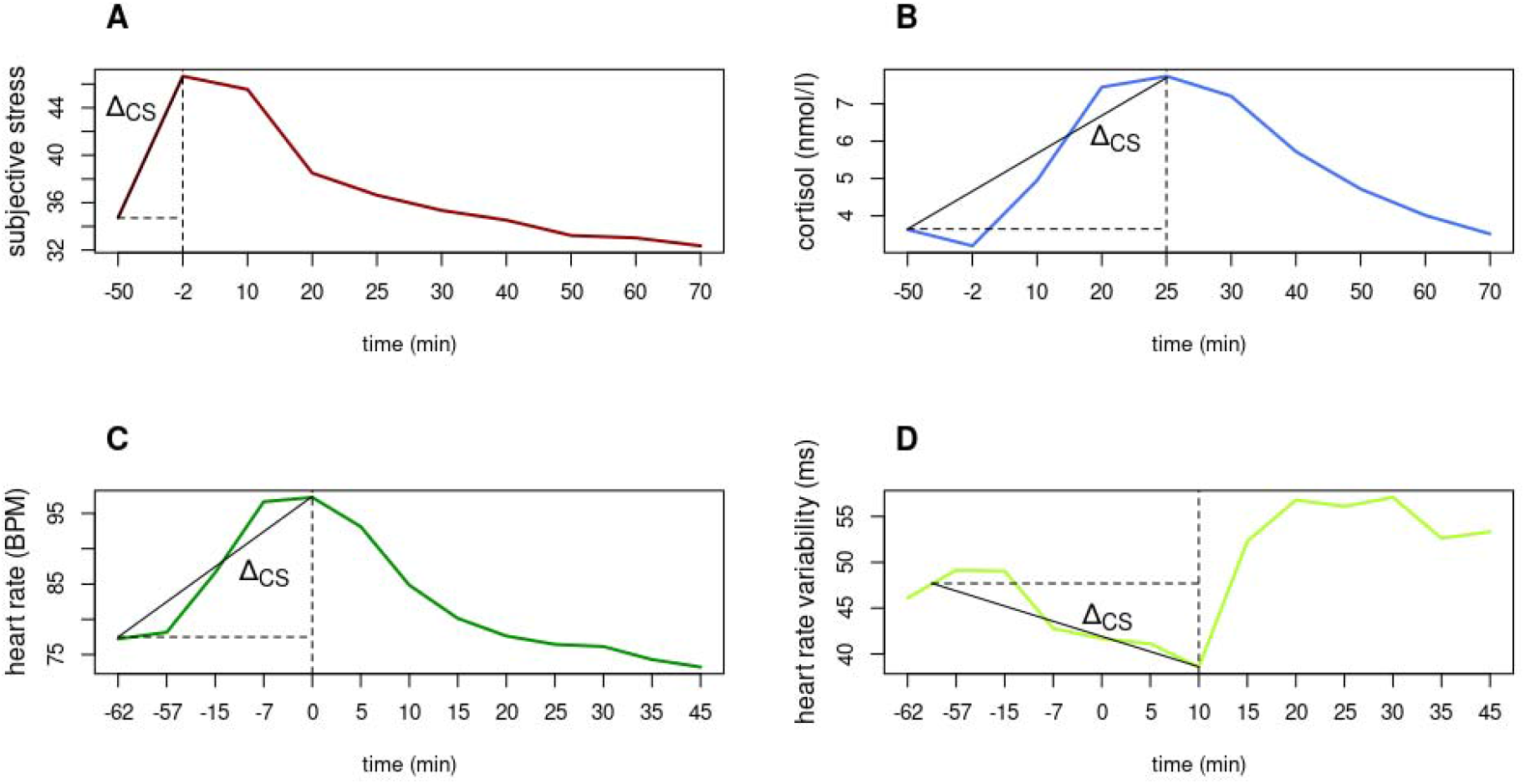
Average raw values of all stress markers. Subjective stress was measured with the STAI state scale. TSST began at 0 min. Time is measured in mins relative to stressor onset. Δ*CS indicates the calculated change scores*.

#### 3.2.1 Association between target relationship variables and target stress

We used linear models predicting target stress reactivity from target-related attachment, dyadic coping and relationship quality. These models revealed a main effect of target insecure-avoidant attachment on subjective stress (β = -15.720, *p* = .004), such that insecure-avoidantly attached individuals reported lower acute subjective stress reactivity (Figure 3). Further, target sex had a significant effect on cortisol reactivity (β = -0.618, *p* = .011), showing the well-established effect of stronger cortisol release in males than females (e.g. Uhart et al., 2006). No other significant effects of target-related predictors on target stress reactivity were found (Table 1).

**Figure 3:**
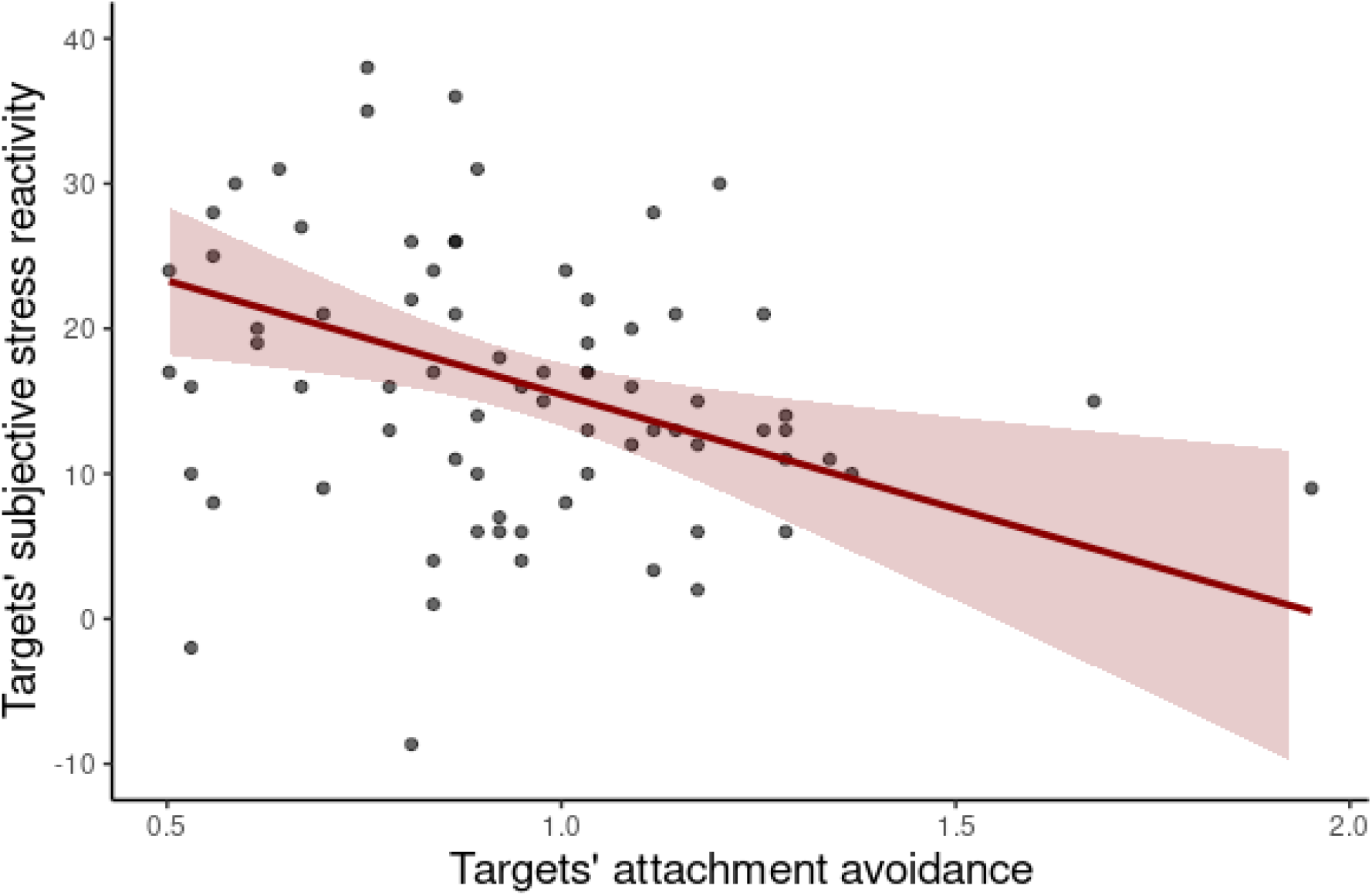
Negative association between target insecure-avoidant attachment and their subjective stress in response to the TSST

**Table 1:** Parameter estimates and fit for target subjective stress, cortisol, heart rate, and HF-HRV reactivity change scores as a function of target variables according to the linear model analyses.

| Target variables → | Subjective stress | Cortisol | Heart rate | Heart rate variability |
| --- | --- | --- | --- | --- |
| ↓ Target variables |  |  |  |  |
| Intercept | **69.371\*\*** | 0.543 | 1.040 | -3.604 |
| Insecure-anxious attachment | 3.943 | -0.511 | -0.429 | 0.295 |
| Insecure-avoidant attachment | **-15.720\*\*** | 0.213 | 0.662 | -0.191 |
| Dyadic coping | -11.550 | 0.110 | 0.078 | 0.983 |
| Relationship quality | -23.062 | 0.546 | 0.417 | 1.422 |
| Age | -8.014 | -0.242 | -0.443 | -0.073 |
| Sex | - | **-0.618\*** | -0.219 | 0.643<sup>a</sup> |
| BMI | - | - | -0.052 | 0.038 |
| Testing start time | - | 0.001 | - | - |
| R<sup>2</sup> | 0.168* | 0.111 | 0.059 | 0.168 |

*Note.* \*\*\*: $p \leq .0005$ . \*\*: $p \leq .005$ . \*: $p \leq .025$ . <sup>a</sup>: $p \leq .05$ for subjective stress and cortisol. \*\*\*:
$p \leq .00025$ . \*\*: $p \leq .0025$ . \*: $p \leq .0125$ . <sup>a</sup>: $p \leq .025$ for heart rate (variability).

#### 3.2.2 Association between observer relationship variables and target stress

Linear regressions predicting targets’ stress reactivity from observers’ attachment, dyadic coping and relationship quality were calculated for each stress marker. Targets showed significantly increased heart rate reactivity with higher insecure-avoidant attachment scores of the observing partners (β = 1.309, *p* = .012; Figure 4). Thus, targets’ heart rate was higher if their observing partner scored higher in insecure-avoidant attachment. Besides the known sex effect on cortisol (β = -0.563, *p = .*016), no other significant effects were found (Table 2).

**Figure 4:**
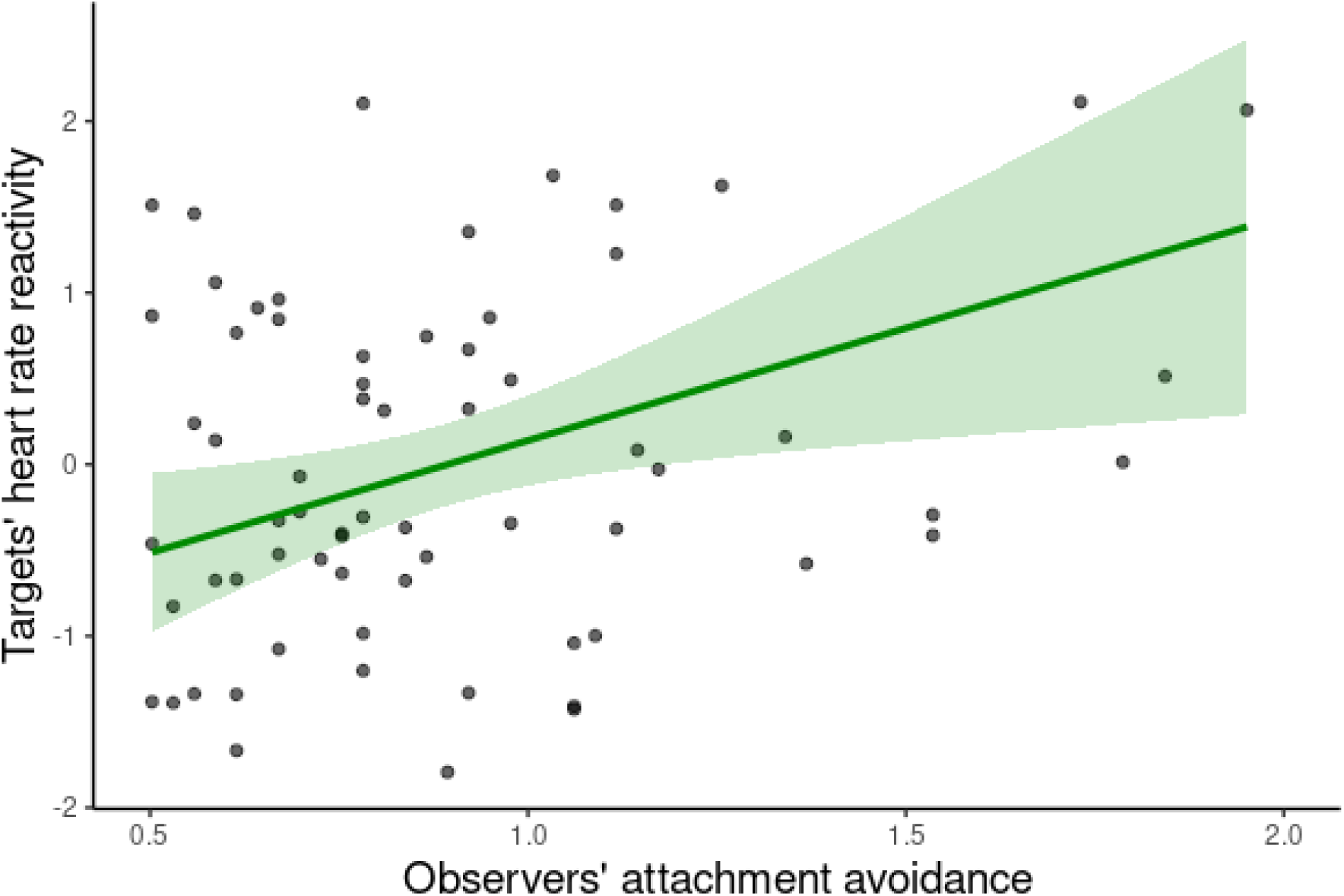
Positive association between observers’ insecure-avoidant attachment scores and target heart rate in response to the TSST

**Table 2:** Parameter estimates and fit for target subjective stress, cortisol, heart rate, and heart rate variability reactivity change scores as a function of observer variables according to the linear model analyses.

| Target variables → | Subjective stress | Cortisol | Heart rate | Heart rate variability |
| --- | --- | --- | --- | --- |
| ↓ Observer variables |  |  |  |  |
| Intercept | **54.248\*** | -0.393 | -1.140 | 0.149 |
| Insecure-anxious attachment | -0.603 | -0.034 | -0.533 | -0.031 |
| Insecure-avoidant attachment | -1.970 | 0.345 | **1.309\*** | -1.124<sup>a</sup> |
| Dyadic coping | -24.873 | 0.495 | 1.427 | 1.004 |
| Relationship quality | 0.463 | 0.378 | 0.535 | -1.114 |
| ↓ Target variables |  |  |  |  |
| Age | -11.067 | -0.384 | -0.708 | 0.118 |
| Sex | - | **-0.563\*** | -0.160 | 0.637 |
| BMI | - | - | -0.032 | 0.024 |
| Testing start time | - | 0.001 | - | - |
| R<sup>2</sup> | 0.092 | 0.091 | 0.145 | 0.250<sup>a</sup> |

*Note.* \*\*\*: $p \leq .0005$ . \*\*: $p \leq .005$ . \*: $p \leq .025$ . a: $p \leq .05$ for subjective stress and cortisol. \*\*\*: $p \leq .00025$ . \*\*: $p \leq .0025$ . \*: $p \leq .0125$ . <sup>a</sup>: $p \leq .025$ for heart rate (variability).

### 3.3 Additional Analyses

A linear regression predicting psychoendocrine covariance from targets’ and observers’ insecure-anxious and -avoidant attachment scores showed a main effect of insecure-avoidant attachment on psychoendocrine covariance (β = -0.302, *p* = .019; Table 3). In detail, individuals with higher insecure-avoidant attachment scores showed significantly less psychoendocrine covariance, meaning a lower correlation between subjective-psychological and cortisol reactivity (Figure 5).

**Figure 5:**
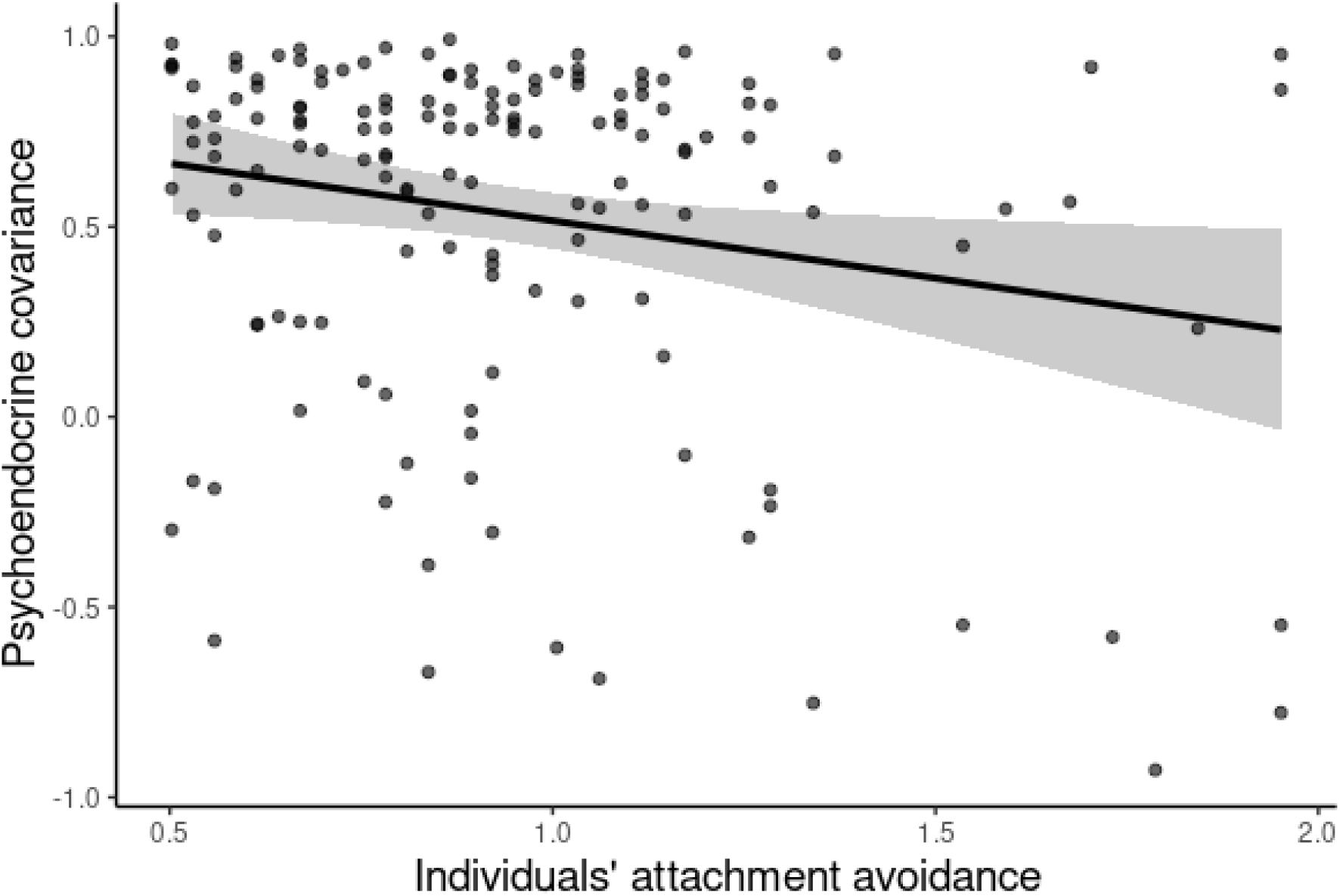
Negative association between participants’ insecure-avoidant attachment scores and their psychoendocrine covariance.

**Table 3:** Parameter estimates and fit for psychoendocrine covariance according to the linear model analyses.

|  | Psychoendocrine covariance |
| --- | --- |
| Intercept | **0.855\*\*** |
| Insecure-anxious attachment | -0.121 |
| Insecure-avoidant attachment | **-0.302\*** |
| Age | 0.029 |
| Sex | -0.059 |
| Testing starting time | 0.000 |
| R<sup>2</sup> | 0.076* |

*Note.* \*\*\*: $p \leq .001$ . \*\*: $p \leq .01$ . \* : $p \leq .05$ . <sup>a</sup>: $p \leq .1$ .

## 4. Discussion

In the current study, we investigated the link between relationship variables and stress reactivity during a psychosocial stress test. During the test, the romantic partner was present but only observed passively. Specifically, we focused on the association between attachment style, relationship quality, and dyadic coping with subjective and physiological stress reactivity during an empathic psychosocial stress paradigm, the empathic Trier Social Stress Test (TSST; Engert et al., 2014a; Kirschbaum et al., 1993). Seventy-nine couples participated in the study, with one partner (the "target") undergoing the TSST and the other partner (the "observer") passively witnessing the situation (Engert et al., 2014b).

Besides focusing on targets’ own relationship variables, this study is first to also consider the association between observers’ relationship variables and targets’ stress reactivity. Insecure-avoidant attachment scores of both targets and observers were found to be significantly linked to targets’ stress. More precisely, targets’ insecure-avoidant attachment scores were negatively correlated with their own subjective stress experience, while observers’ insecure-avoidant attachment scores were positively correlated with targets’ heart rate reactivity. Further, higher insecure-avoidant attachment scores in both targets and observers were associated with lower psychoendocrine covariance, that is, a lower accordance between self-reported and cortisol stress responding. Our study therefore highlights the role of individual differences in one’s own and the romantic partner’s insecure-avoidant attachment in the experience of acute stressors.

The finding of lower subjective stress reports in targets with higher insecure-avoidant attachment scores confirms our a priori hypothesis, and is in line with previous studies employing relationship-related stressors rather than an “external” psychosocial paradigm such as the TSST (Dewitte et al., 2010; Diamond et al., 2006). Although target insecure-avoidant attachment scores were not linked to physiological stress reactivity, they were linked to lower psycho-endocrine covariance. This may suggest a reduced ability to recognize or report one’s own stress levels. To that effect, rather than actually reducing stress, higher insecure-avoidant attachment may increase the use of emotion suppression. This interpretation is in line with attachment theory, which suggests that individuals with higher insecure-avoidant attachment are more likely to employ deactivating attachment strategies, besides being more prone to rely on emotional suppression as a coping mechanism to preserve their sense of independence (Simpson & Rholes, 2017).

Intriguingly, and confirming our hypothesis, targets’ stress reactivity was also associated with partners’ insecure-avoidant attachment scores, such that stress-induced increases in targets’ heart rate were stronger when the observing partner scored higher on insecure-avoidant attachment. Thus, when confronted with a non-relational stressor, next to one’s own attachment style, the attachment style of one’s romantic partner may also contribute to the intensity of one’s stress response. This finding extends the work by Dewitte et al. (2010), who found that individuals with their securely attached partners present during a relational distress paradigm had the lowest stress responses (Dewitte et al., 2010). It further aligns with a study using a relationship discussion stressor (Brooks et al., 2011) in which women responded with higher cortisol release if they were discussing with an insecure-avoidantly attached partner. From an attachment theory perspective, the insecure-avoidant observers’ tendency to deactivate their attachment system, in ways of suppressing their own feelings and shutting out others’ vulnerability, may pressure targets to hide their feelings, which may in turn aggravate the intensity of the experienced stressor.

Whether or not insecure-avoidantly attached individuals actually suppressed their stress, felt less stress, or just reported to do so, we cannot answer conclusively in the context of our study. However, when considering the perspective of the observing partner, insecure-avoidant target attachment does not seem to be the most beneficial prerequisite for social support during stress. Therefore, modifying behavioral patterns typically associated with insecure-avoidant attachment, such as emotional suppression or withdrawal, may be beneficial for dyadic stress reduction.

We found no associations of targets’ stress with insecure-anxious attachment, dyadic coping and relationship quality of either targets or observers. The lack of an association between targets’ stress reactivity and their own as well as their partners’ insecure-anxious attachment could be explained by the fact that one of the main components of insecure-anxious attachment is a fear of abandonment (Simpson & Rholes, 2017). Hence, because observers were always present during the TSST as they could not leave the room, we may speculate that their continuous presence was unlikely to trigger such a fear of abandonment in targets. Furthermore, because insecure-anxiously attached individuals are more likely to engage in heightened, more intensive caregiving – rather than deactivating caregiving as associated with insecure-avoidant attachment – (Colledani et al., 2022), it may be that the insecure-anxious attachment dimension in observers also had less of an influence on targets’ stress response. The lack of predictive power of relationship quality and dyadic coping may be explained by the fact that despite being passively present, observing partners had little power to directly influence the unfolding TSST, and were un available for joint stress reduction afterwards. Thus, the protective properties of relationship quality or dyadic coping ability may not have been as pronounced as otherwise possible.

Not all stress markers were equally affected. Specifically, targets’ insecure-avoidant attachment appeared to be linked to their subjective stress, which we interpret as an indicator for an attachment system deactivation, possibly through emotional suppression and an emphasis on independence. What was measured might therefore be a coping strategy rather than a direct impact on stress reactivity. Additionally, while observer’s insecure-avoidant attachment was unrelated to their stress levels as measured by the main stress hormone cortisol, it did associate with heart rate. Targets’ elevated heart rate may be indicative of increased agitation or arousal (rather than actual stress), potentially due to a heightened state of vigilance in the target. Therefore, it appears that different physiological markers may reflect attachment processes in different ways. This highlights the nuanced and complex interaction between relationship variables and stress responding, suggesting that a multifaceted approach is necessary to fully capture these effects.

There are some limitations to this study. First, variance in attachment scores was low, likely due to our restrictive inclusion criteria that may have led to a sample with low levels of insecure-avoidant and -anxious attachment. Additionally, because we did not directly assess emotional suppression, we cannot conclusively determine whether individuals with higher insecure-avoidant attachment actually suppressed their feelings, experienced less stress, or merely reported to have done so.

In sum, we here examined the association between relationship variables, including attachment style, of both targets and observers with the targets’ stress experience during a non-relational stressor. We found insecure-avoidant attachment of targets and observers to correlate with targets’ stress reactivity, such that their own insecure-avoidant attachment scores were associated with their subjective stress experience, while observers’ insecure-avoidant attachment scores were associated with heart rate reactivity in the target. Further, insecure-avoidant attachment of both targets and observers was linked to their psycho-endocrine covariance, indicating potential suppression of emotional experiences. We conclude that the presence of an insecure-avoidantly attached partner during stressful experiences may increase an individual’s physiological stress reactivity, rather than being a source of support. Long-term, this may negatively impact stress- related health and wellbeing.

## Acknowledgements

We are thankful to the members of the “Social Stress and Family Health” Research Group at the MPI CBS involved in the ARC project, particularly Elisabeth Murzik, Michael Vollmann, and Sylvia Tydecks, for managing the study, to Henrik Grunert for technical assistance, and the many student assistants and interns for recruitment, data collection, and data processing.

