## SupplementaryMaterial for "Relationship as resource or burden? Associations of attachment style, relationship quality and dyadic coping with acute psychosocial stress in the presence of the romantic partner"

Corresponding author:

Mathilde Gallistl

**Table S1**

*Means, standard deviations and correlations for the considered predictors in the target and observer.*

| Variable | M | SD | 1 | 2 | 3 | 4 | 5 | 6 | 7 | 8 | 9 | 10 | 11 |
| --- | --- | --- | --- | --- | --- | --- | --- | --- | --- | --- | --- | --- | --- |
| Target Variables | | | | | | | | | | | | | |
| 1. Age | 26.22 | 4.41 |  |  |  |  |  |  |  |  |  |  |  |
| 2. Sex | 0.49 | 0.50 | -.09 |  |  |  |  |  |  |  |  |  |  |
| 3. BMI | 23.01 | 3.05 | .03 | -.25* |  |  |  |  |  |  |  |  |  |
| 4. Start time of testing | 181.76 | 75.42 | -.07 | -.01 | -.05 |  |  |  |  |  |  |  |  |
| 5. Attachment Anxiety | 2.38 | 0.77 | .12 | -.13 | .12 | .04 |  |  |  |  |  |  |  |
| 6. Attachment Avoidance | 1.88 | 0.55 | .31** | -.14 | -.02 | .13 | .41** |  |  |  |  |  |  |
| 7. Relationship Quality | 30.45 | 3.44 | -.12 | .07 | -.08 | .10 | -.29** | -.51** |  |  |  |  |  |
| 8. Dyadic Coping | 136.56 | 13.41 | -.27* | .22 | -.03 | .01 | -.27* | -.60** | .51** |  |  |  |  |
| Observer Variables | | | | | | | | | | | | | |
| 9. Attachment Anxiety | 2.32 | 0.77 | .15 | -.05 | .01 | -.06 | .24* | .25* | -.48** | -.24* |  |  |  |
| 10. Attachment Avoidance | 1.86 | 0.76 | .25* | -.08 | -.11 | -.10 | .26* | .30** | -.41** | -.33** | .50** |  |  |
| 11. Relationship Quality | 30.74 | 3.83 | -.19 | .03 | .04 | .06 | -.25* | -.27* | .45** | .41** | -.50** | -.70** |  |
| 12. Dyadic Coping | 141.37 | 15.16 | -.10 | .01 | -.04 | .22 | -.20 | -.26* | .46** | .46** | -.35** | -.56** | .64** |

*Note*. Participants’ sex was coded as “0” male, “1” female. Start time of testing is coded in minutes after 12pm. Values for attachment related avoidance and anxiety, relationship quality, and dyadic coping were winsorized. M and SD are used to represent mean and standard deviation, respectively.

* indicates p < .05. ** indicates p < .01.
